# Regulation of *X. laevis* M18BP1 centromeric localization and CENP-A assembly

**DOI:** 10.1101/2025.07.15.664882

**Authors:** Rae R. Brown, Jacob P. Schwartz, Lyin Ghadri, Aaron F. Straight

**Author notes:** Correspondence, 279 Campus Drive, B409A Stanford, CA 94305.

## Abstract

Eukaryotic chromosome segregation requires attachment of chromosomes to microtubules of the mitotic spindle through the kinetochore so that chromosomes can align and move in mitosis. Kinetochores are assembled on the centromere which is a unique chromatin domain that is epigenetically defined by the histone H3 variant CENtromere Protein A (CENP-A). During DNA replication CENP-A is equally divided between replicated chromatids and new CENP-A nucleosomes are re-assembled during the subsequent G1 phase of the cell cycle. How cells regulate the strict cell cycle timing of CENP-A assembly is a central question in the epigenetic maintenance of centromeres and kinetochores. One essential assembly factor for CENP-A nucleosomes is the Mis18 complex (Mis18α, Mis18β, and M18BP1) which is regulated in its localization to centromeres between metaphase and G1 when CENP-A assembly occurs. Here, we define a new regulatory mechanism that works through cell cycle dependent phosphorylation of *Xenopus laevis* M18BP1 between metaphase and interphase. This phosphoregulatory switch disrupts binding of M18BP1 to CENP-A nucleosomes in metaphase, and when relieved enables M18BP1 binding to CENP-A nucleosomes in interphase. We show that this phosphorylation dependent switching mechanism regulates CENP-A nucleosome assembly. We propose that the phospho-regulated binding of M18BP1 to CENP-A nucleosomes is an important control mechanism that restricts the timing of new CENP-A assembly.

## Introduction

All eukaryotic cells undergo the fundamental process of cell division, wherein each daughter cell receives an equal chromosome complement. Central to this process is the centromere, the region of the chromosome to which the mitotic spindle attaches for accurate chromosome segregation. The centromere is epigenetically defined by the histone H3 variant CENtromere Protein A (CENP-A) (Warburton et al. 1997; Palmer et al. 1987; Sullivan, Hechenberger, and Masri 1994; Earnshaw and Rothfield 1985; Palmer et al. 1991; Meluh et al. 1998; Takahashi, Chen, and Yanagida 2000; Henikoff et al. 2000). Loss of centromere identity and disruption of CENP-A results in aneuploidy, chromosome instability, and cell death (Stoler et al. 1995; Blower and Karpen 2001; Howman et al. 2000; Régnier et al. 2005; Goshima et al. 2003).

With each round of cell division, CENP-A nucleosomes are split between daughter cells, and new CENP-A nucleosomes must be assembled in order to faithfully maintain the centromere. H3 nucleosomes are assembled during the S phase of the cell cycle and coupled with DNA replication (Groth et al. 2007; Ramachandran and Henikoff 2015; Worcel, Han, and Wong 1978). However, in many metazoans, CENP-A nucleosome assembly is decoupled from DNA replication, and occurs during late telophase/early G1 (Jansen et al. 2007; Maddox et al. 2007; Moree et al. 2011; Bernad et al. 2011; Silva et al. 2012). Although proteins involved in this process have been identified, it is unknown exactly how this assembly timing is controlled.

Holliday JUnction Recognition Protein (HJURP) is a histone chaperone for the CENP-A:H4 dimer that is necessary for CENP-A nucleosome assembly. However, HJURP cannot localize to the centromere and bind to CENP-A nucleosomes on its own (Dunleavy et al. 2009; Foltz et al. 2009). In frogs (*Xenopus laevis*) and chickens (*Gallus gallus*), both the Mis18 complex (a hetero-octameric complex composed of Mis18α tetramer, a Mis18β dimer, and an M18BP1 dimer) and CENtromere Protein C (CENP-C) bind directly to CENP-A nucleosomes. Each of these proteins binds to HJURP, thereby serving as adaptors to localize HJURP:CENP-A:H4 to centromeres and facilitate the deposition of the CENP-A:H4 histone dimer. However, while CENP-C is constitutively localized to the centromere, the Mis18 complex only localizes in interphase/G1 (Moree et al. 2011; French et al. 2017; Carroll, Milks, and Straight 2010; Watanabe et al. 2019; Ariyoshi et al. 2021; Jiang et al. 2023). In humans, the Mis18 complex cannot bind to CENP-A nucleosomes. Instead, the Mis18 complex binds to CENP-C to facilitate CENP-A assembly (McKinley and Cheeseman 2014; Nardi et al. 2016; Fujita et al. 2007; Maddox et al. 2007; Pan et al. 2019; Moree et al. 2011; Dambacher et al. 2012).

Understanding how the Mis18 complex directly engages CENP-A nucleosomes and how that interaction changes between metaphase and interphase is important for understanding the cell cycle regulation of CENP-A assembly. *X. laevis* and *G. gallus* M18BP1 contain a CENP-C motif which is absent in humans. This motif enables binding to CENP-A nucleosomes via the same mechanism used by CENP-C (Hori et al. 2017; French et al. 2017; Kral 2015; Jiang et al. 2023). *X. laevis* is allotetraploid with two subgenomes, and as such it has two isoforms of M18BP1: M18BP1-S (1) and M18BP1-L (2). Both isoforms have the conserved CENP-C motif and bind to CENP-A nucleosomes *in vitro* (French et al. 2017). In interphase egg extract, both M18BP1-L and M18BP1-S bind to CENP-A nucleosomes as part of the Mis18 complex to facilitate CENP-A assembly. However, in metaphase egg extract, the Mis18 complex dissociates and neither M18BP1 isoform binds to CENP-A nucleosomes. It remains unclear what disrupts this interaction such that M18BP1 is no longer bound to CENP-A nucleosomes in metaphase.

CENP-C, the Mis18 complex, and HJURP are regulated by mitotic kinases, and in particular Polo-like Kinase 1 (Plk1) and Cyclin-dependent kinase 1 (Cdk1) (McKinley and Cheeseman 2014; Silva et al. 2012; Pan et al. 2017; French and Straight 2019; Stankovic et al. 2017; Spiller et al. 2017; Müller et al. 2014). Cdk1 phosphorylation triggers the disassembly of the Mis18 complex and disrupts HJURP localization to centromeres by blocking its Mis18 complex and CENP-C binding (McKinley and Cheeseman 2014; Pan et al. 2017; Spiller et al. 2017; Müller et al. 2014; French et al. 2017; Pan et al. 2019; French and Straight 2019; Stankovic et al. 2017; Wang et al. 2014; Stellfox et al. 2016).

In this work, we examine the cell cycle regulation of CENP-A assembly in *X. laevis*. We find that the mitotic phosphorylation of a conserved residue, serine 760 (S760), in the CENP-C motif of M18BP1-L prevents binding to CENP-A nucleosomes, thereby restricting CENP-A assembly to interphase. Consistent with this, phosphomimetic mutation of the S760 residue of M18BP1-L to aspartic acid (S760D) in interphase extracts, when CENP-A is normally assembled, disrupts its localization to centromeres and inhibits CENP-A assembly. In metaphase egg extract, a non phosphorylatable mutant of M18BP1-L, S760 mutated to alanine (S760A), is unable to localize to the centromere, suggesting additional mechanism(s) preventing metaphase binding of M18BP1-L binding to CENP-A nucleosomes.

## Results

### M18BP1-L binding to CENP-A nucleosomes is disrupted by a conserved mitotic phosphorylation

To identify residues in M18BP1-L that regulate the timing of its centromere localization, we previously mapped mitotic phosphorylation sites in its CENP-A binding domain. Of these sites, the Cdk1 consensus phosphorylations did not disrupt the binding of M18BP1-L to CENP-A nucleosomes *in vitro* (French et al. 2017). Thus, we focused our analysis on the next most highly phosphorylated residue, serine 760. Interestingly, this serine is located near the conserved arginine anchor identified as being important for the affinity of M18BP1^ggKNL2^ for CENP-A nucleosomes via the H2A/H2B acidic patch (Jiang et al. 2023). It is similarly conserved across species that have the conserved CENP-C motif in M18BP1 (Fig 1A) (Kral 2015; Jiang et al. 2023; French et al. 2017). We hypothesized that this phosphorylation could disrupt the association between M18BP1-L and CENP-A nucleosomes.

**Figure 1.**
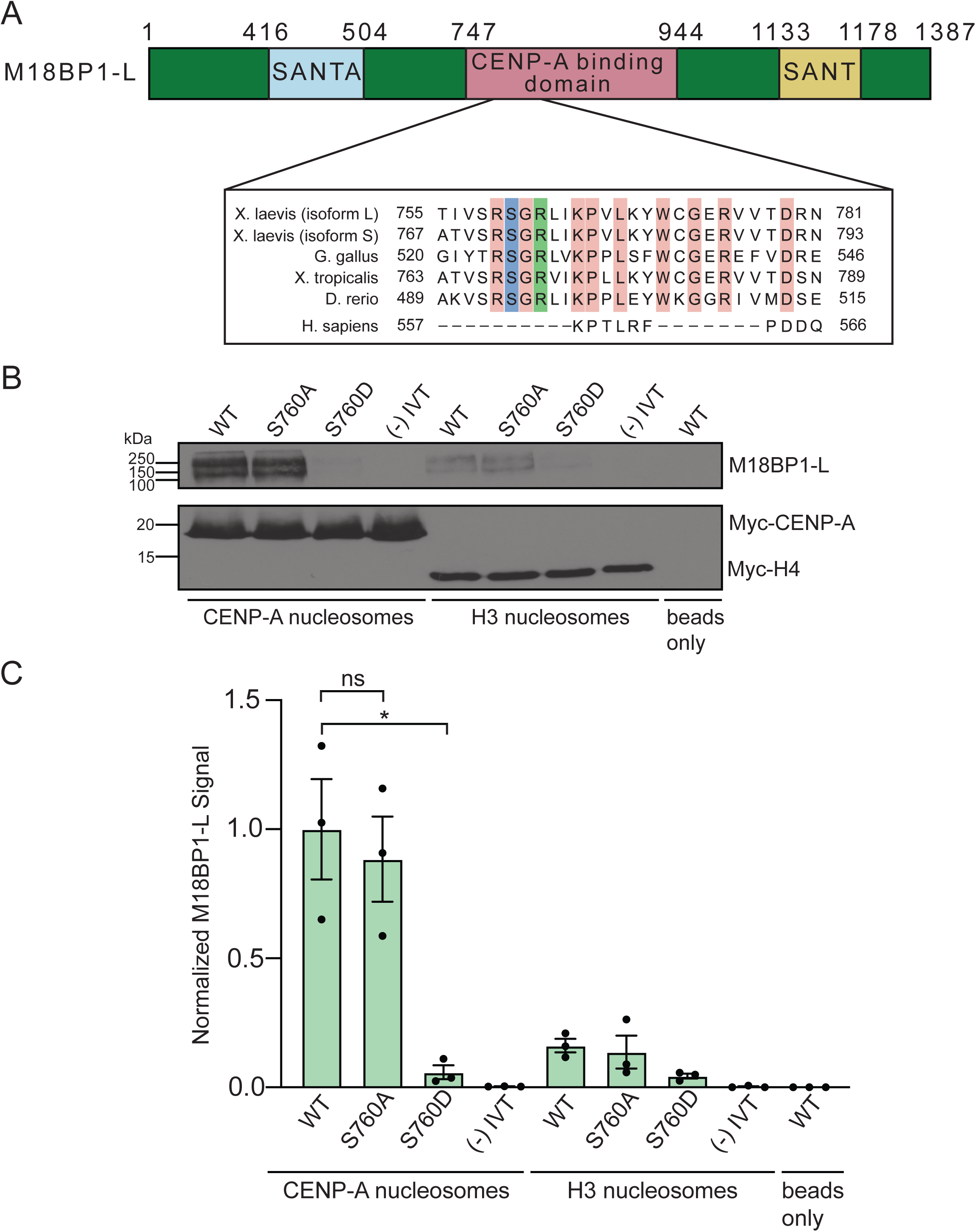
**A**. Diagram of *X. laevis* M18BP1-L protein. The domain required for CENP-A nucleosome binding is shown in pink and the SANT and SANTA domains are highlighted in yellow and blue respectively. The inset shows the region homologous to the CENP-C binding motif, with sequence alignment for *X. laevis* (both isoforms), *G. gallus*, *X. tropicalis*, *D. rerio*, and *H. sapiens*. 100% conserved residues for all species except human are highlighted in pink, the conserved arginine described in (Jiang et al. 2023) is highlighted in green, and the conserved serine shown in this work is highlighted in blue. **B**. Western blot assay of M18BP1 binding to CENP-A or H3 nucleosome arrays. M18BP1 WT, phosphorylation site mutants (S760A and S760D), and a negative control (-) IVT are indicated. The top panel displays an anti-xlM18BP1 western blot and the bottom panel displays an anti-myc blot to control for chromatin levels. The last lane displays a bead only (no nucleosome) negative control. **C**. Quantification of western blots of M18BP1 binding to CENP-A or H3 nucleosomes. The amount of protein bound to chromatin is normalized to the WT M18BP1 binding to CENP-A nucleosomes. Error bars represent SEM of three independent replicates (n = 3). \**P = 0.0379, ^ns^P = 0.6744*.

To test the role of S760 in CENP-A nucleosome binding, we mutated serine 760 to alanine (S760A) or to aspartic acid (S760D) to prevent or to mimic phosphorylation respectively. We expressed these proteins and wild type (WT) M18BP1-L using *in vitro* transcription and translation. We then assayed the binding of WT M18BP1-L, M18BP1-L^S760A^, and M18BP1-L^S760D^ to CENP-A nucleosomes by reconstituting CENP-A nucleosomes *in vitro* on an 18x array of the 601 Widom sequence (Lowary and Widom 1998) and western blotting the bound protein. WT M18BP1-L and M18BP1-L^S760A^ bind to CENP-A nucleosomes *in vitro* with similar levels, whereas M18BP1-L^S760D^ fails to bind to CENP-A nucleosomes (Fig 1B, 1C). These results indicate that the S760 phosphorylation site of M18BP1-L regulates CENP-A nucleosome binding in *X. laevis*.

### M18BP1-L phospho-mimetic mutations prevent interphase centromere localization and reduce CENP-A assembly

Next, we tested if phosphorylation of S760 restricts M18BP1 centromere localization to interphase. We assayed centromeric localization of phospho-null M18BP1-L^S760A^ and phospho-mimetic M18BP1-L^S760D^ on sperm nuclei in interphase *X. laevis* egg extracts. We immunodepleted endogenous M18BP1 (both isoforms) from egg extract and added back *in vitro* transcribed and translated WT M18BP1-L, M18BP1-L^S760A^, or M18BP1-L^S760D^ (Fig 2A, 2B), and then assayed the localization using immunofluorescence. We found that, consistent with the *in vitro* nucleosome binding results, M18BP1-L WT and M18BP1-L^S760A^ localization to the centromere was normal. In contrast, phospho-mimetic M18BP1-L^S760D^ is significantly reduced to ∼40% of WT levels (Fig 2C, 2D). This is consistent with phosphorylation of M18BP1-L S760 inhibiting centromere localization in *X. laevis* egg extract and CENP-A nucleosome binding *in vitro*.

**Figure 2.**
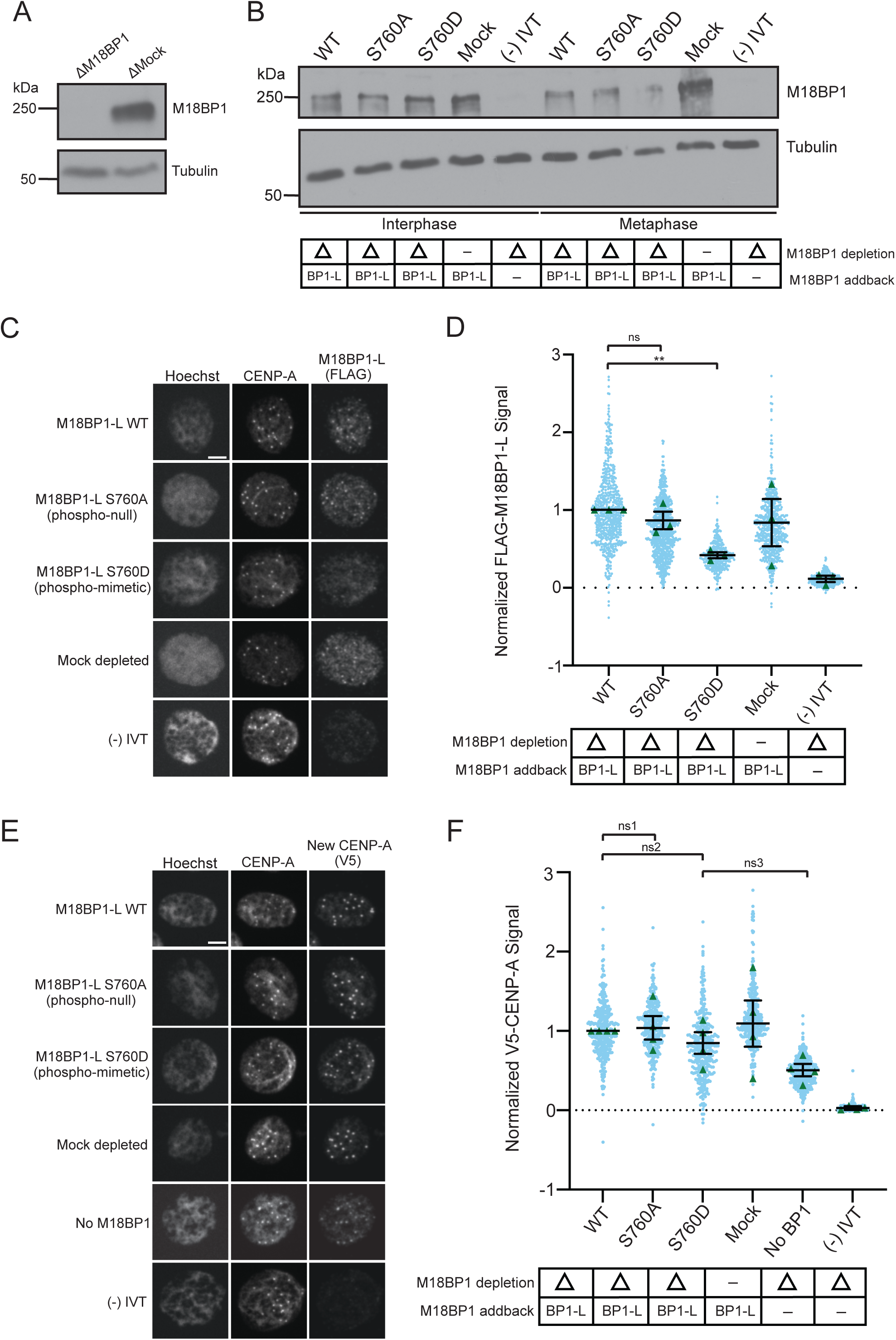
**A**. A representative western blot showing the efficiency of immunodepletion of endogenous M18BP1 from *X. laevis* egg extract. Depletion (Δ) or mock depletion with rabbit IgG is indicated above each column. The M18BP1 blot is shown in the top panel and a tubulin loading control is shown in the bottom panel. **B**. A representative western blot showing the addback of WT or mutant M18BP1 in egg extract. The depletion condition (Δ) is indicated in the table below the blot. The M18BP1 blot is shown in the top panel and the tubulin loading control is shown in the bottom panel. **C**. Representative immunofluorescence images of full-length WT or mutant FLAG-M18BP1-L localization with mock depletion and (-) IVT control (indicated on left) in interphase extract immunodepleted of endogenous M18BP1. Labeling for DNA (Hoechst), total CENP-A, and M18BP1-L (FLAG) is indicated above the image. Scale bar is 5μm. **D**. Quantification of WT or mutant FLAG-M18BP1-L localization with controls (indicated below) in interphase egg extract immunodepleted of endogenous M18BP1. The M18BP1 depletion and addback condition is indicated in the bottom table. The mean signal is normalized to WT FLAG-M18BP1-L localization. Error bars represent SEM of three independent replicates (n = 3) with green triangles displaying the mean of each replicate and blue circles representing each individual centromere. *\*\*P = 0.0041, ^ns^P = 0.3525*. **E**. Representative immunofluorescence images of new V5-CENP-A assembly in interphase extract immunodepleted of endogenous M18BP1 then supplemented with full-length WT or mutant FLAG-M18BP1-L, a mock depletion with rabbit IgG, a (-) IVT control without V5-CENP-A, or no M18BP1 addback (no BP1) (indicated on left). Labeling for DNA (Hoechst), total CENP-A, and new CENP-A (V5) is indicated above the image. Scale bar is 5μm. **F**. Quantification of new V5-CENP-A with controls (indicated below) in interphase egg extract immunodepleted of endogenous M18BP1. M18BP1 depletion and addback condition indicated in the bottom table. The signal is normalized to the WT FLAG-M18BP1-L addback condition. Error bars represent SEM of four independent replicates (n = 4) with green triangles displaying the mean of each replicate and blue circles representing each individual centromere. *^ns1^P = 0.8053, ^ns2^P = 0.3465, ^ns3^P = 0.0821*.

Given that the phosphomimetic mutant of M18BP1-L has reduced localization at centromeres during interphase, we tested whether it affected CENP-A nucleosome assembly in *X. laevis* egg extract. We immunodepleted endogenous M18BP1 from egg extract, then added *in vitro* transcribed and translated WT M18BP1-L, M18BP1-L^S760A^, or M18BP1-L^S760D^. We supplemented the extract with V5-CENP-A mRNA to track new CENP-A assembly via the V5 tag. CENP-A assembly is lower in M18BP1-L^S760D^, although it is not statistically significantly different from WT M18BP1-L and M18BP1-L^S760A^ (p=0.3525) (Fig 2E, 2F). Extracts that were not complemented with M18BP1 (no BP1) showed some CENP-A assembly compared to the control without added V5-CENP-A ((-) IVT) indicating residual assembly activity in the absence of M18BP1 (Fig 2E, 2F).

### Removal of CENP-C uncovers a role for M18BP1-L phosphorylation in CENP-A assembly

We previously showed that depletion of CENP-C leads to an increase in M18BP1 localization to the centromere and that full CENP-A assembly requires both CENP-C and M18BP1 (French et al. 2017). Therefore, we tested whether the presence of CENP-C might impact the S760 phosphorylation dependent localization of M18BP1-L. To do this, we immunodepleted both endogenous M18BP1 and CENP-C and then complemented the extracts with WT or mutant M18BP1-L (Fig 3A). Consistent with previous results, the levels of WT M18BP1-L at centromeres were higher than the mock depletion condition, in which CENP-C is still present (Fig 3B, 3C). The phospho-null mutant M18BP1-L^S760A^ localized to WT levels. The phospho-mimetic M18BP1-L^S760D^ had reduced localization in the absence of CENP-C, to ∼30% WT levels. Thus, in the presence or absence of CENP-C, phosphomimetic M18BP1-L is reduced in its levels at centromeres.

**Figure 3.**
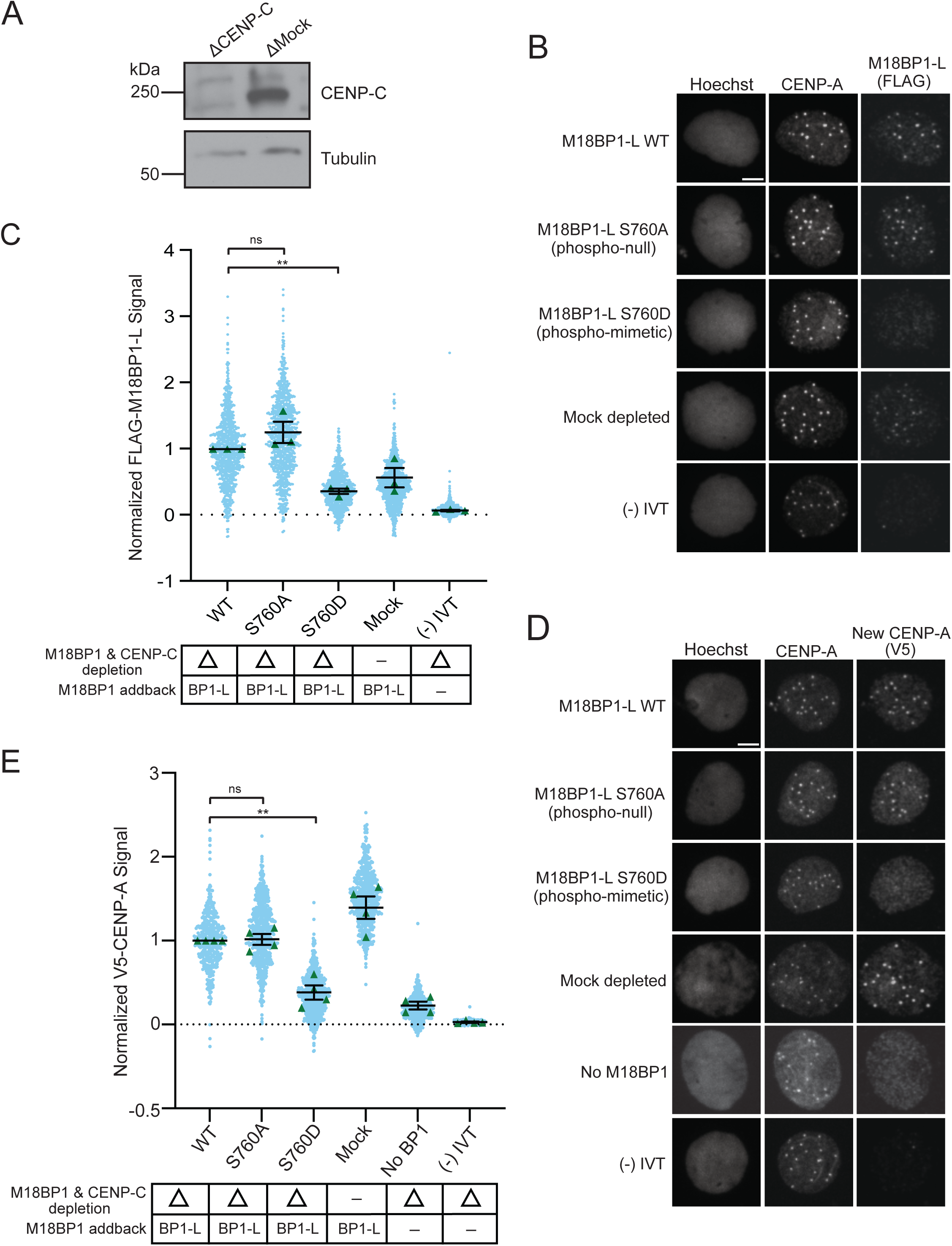
**A**. A representative western blot displaying immunodepletion of endogenous CENP-C and an IgG mock depletion using in egg extract. The depletion condition (Δ) is indicated above each column. The CENP-C blot is shown in the top panel and a tubulin loading control is shown in the bottom panel. **B**. Representative immunofluorescence images of full-length WT or mutant FLAG-M18BP1-L localization with mock depletion and (-) IVT control (indicated on left) in interphase extract immunodepleted of endogenous CENP-C and M18BP1. Labeling for DNA (Hoechst), total CENP-A, and M18BP1-L (FLAG) is indicated above the image. Scale bar is 5μm. **C**. Quantification of WT or mutant FLAG-M18BP1-L localization with controls (indicated below) in interphase egg extract immunodepleted of endogenous CENP-C and M18BP1. CENP-C and M18BP1 depletion and addback conditions are indicated in the bottom table. The signal is normalized to WT FLAG-M18BP1-L localization. Error bars represent SEM of three independent replicates (n = 3) with green triangles displaying the mean of each replicate and blue circles representing each individual centromere. *\*\*P = 0.0041, ^ns^P = 0.2570*. **D**. Representative immunofluorescence images of new V5-CENP-A assembly in interphase extract immunodepleted of endogenous CENP-C and M18BP1 then supplemented with full-length WT or mutant FLAG-M18BP1-L, a mock depletion, a (-) IVT control without V5-CENP-A, or no M18BP1 addback (no BP1) (indicated on left). Labeling for DNA (Hoechst), total CENP-A, and new CENP-A (V5) is indicated above the image. Scale bar is 5μm. **E**. Quantification of new V5-CENP-A with controls (indicated below) in interphase egg extract immunodepleted of endogenous CENP-C and M18BP1. CENP-C and M18BP1 depletion and addback condition indicated in the bottom table. The signal is normalized to the WT FLAG-M18BP1-L addback condition. Error bars represent SEM of four independent replicates (n = 4) with green triangles displaying the mean of each replicate and blue circles representing each individual centromere. *\*\*P = 0.0056, ^ns^P = 0.8257*.

Because CENP-C and M18BP1 are required for CENP-A assembly in *X. laevis* (French et al. 2017), we tested if the absence of CENP-C would exacerbate the effect of M18BP1-L S760 phosphorylation on CENP-A assembly. We repeated the experiment with immunodepletion of both endogenous M18BP1 and CENP-C from egg extract followed by complementation with M18BP1-L and V5-tagged CENP-A. In the absence of CENP-C, new CENP-A assembly is significantly reduced with the addition of M18BP1-L^S760D^ compared to the addition of WT M18BP1-L or M18BP1-L^S760A^ or mock depleted extract (Fig 3D, 3E). This data shows that phosphorylation of M18BP1-L controls its proper localization to centromeres and CENP-A assembly, and that both M18BP1-L and CENP-C contribute to proper CENP-A assembly.

### Dephosphorylation of M18BP1-L is not sufficient for bypass of metaphase inhibition of centromere localization or CENP-A assembly

M18BP1-L does not localize to the centromere in metaphase when S760 is phosphorylated, unlike its isoform M18BP1-S (French and Straight 2019; Flores Servin, Brown, and Straight 2023). We tested whether lack of S760 phosphorylation would be sufficient to cause metaphase centromere localization and CENP-A assembly by immunodepleting endogenous M18BP1 from metaphase extracts, adding back WT M18BP1-L, M18BP1-L^S760A^, or M18BP1-L^S760D^, and then assaying M18BP1-L localization to sperm nuclei. WT M18BP1-L, M18BP1-L^S760A^, and M18BP1-L^S760D^ all failed to localize to the centromere in metaphase egg extract (Fig 4A, 4B). CENP-C localizes to the centromere throughout the cell cycle, and could be competing with M18BP1-L^S760A^ binding to CENP-A nucleosomes in metaphase. To test this, we immunodepleted both CENP-C and M18BP1, and assayed WT M18BP1-L, M18BP1-L^S760A^, and M18BP1-L^S760D^ localization. Similar to conditions where CENP-C is present, in the absence of CENP-C M18BP1-L fails to localize to centromeres regardless of S760 mutation (Fig 4C, 4D). Additionally, CENP-A assembly does not occur in metaphase in all phospho-mutant and immunodepletion conditions (Fig EV1A – EV1D). This indicates that additional activities prevent M18BP1-L binding to CENP-A nucleosomes and CENP-A assembly in metaphase.

**Figure 4.**
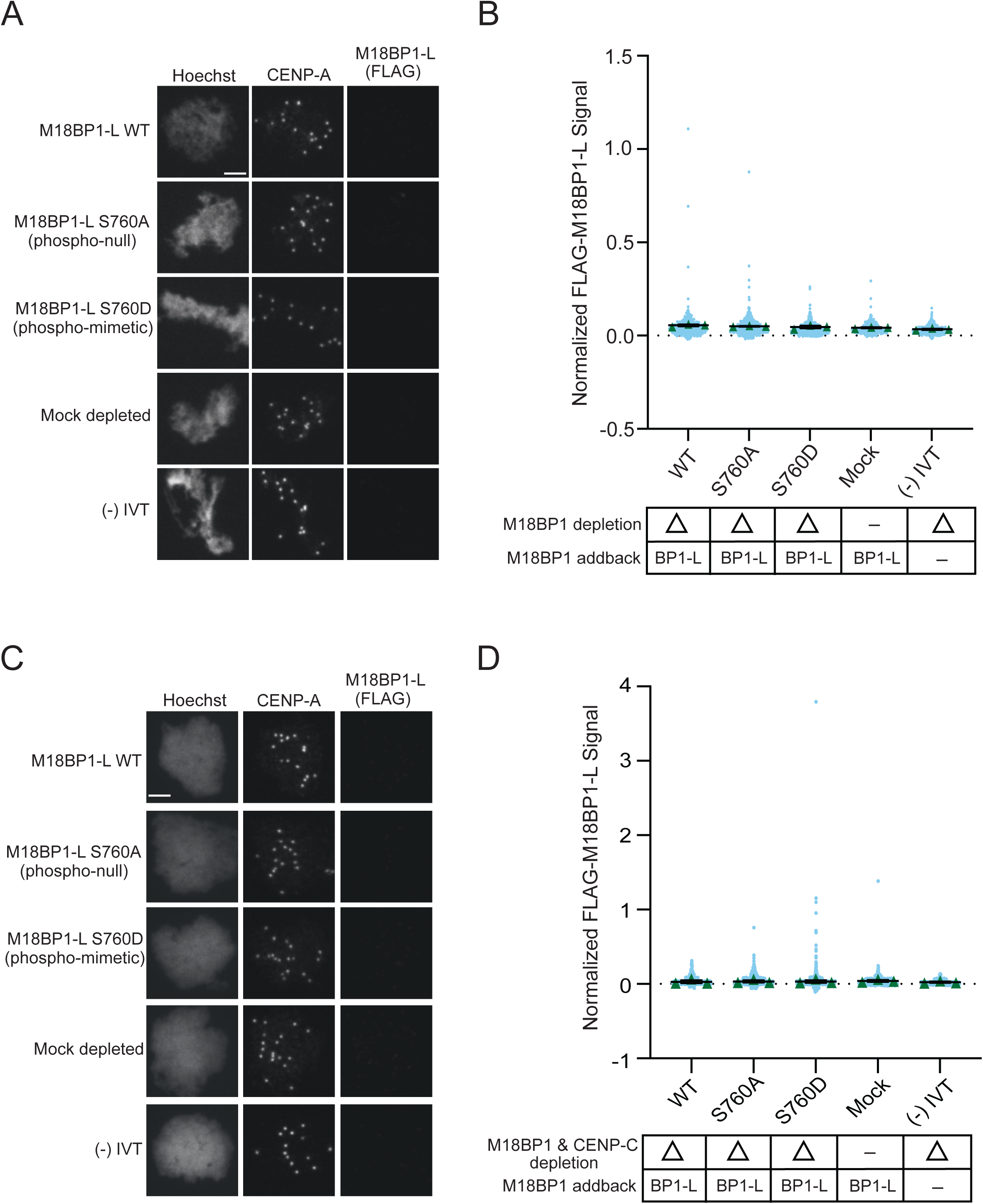
**A**. Representative immunofluorescence images of full-length WT or mutant FLAG-M18BP1-L localization with mock depletion and (-) IVT control (indicated on left) in metaphase extract immunodepleted of endogenous M18BP1. Labeling for DNA (Hoechst), total CENP-A, and M18BP1-L (FLAG) is indicated above the image. Scale bar is 5μm. **B**. Quantification of WT or mutant FLAG-M18BP1-L localization with controls (indicated below) in metaphase egg extract immunodepleted of endogenous M18BP1. The M18BP1 depletion and addback condition is indicated in the bottom table. The signal is normalized to WT FLAG-M18BP1-L localization. Error bars represent SEM of three independent replicates (n = 3) with green triangles displaying the mean of each replicate and blue circles representing each individual centromere. **C**. Representative immunofluorescence images of full-length WT or mutant FLAG-M18BP1-L localization with mock depletion and (-) IVT control (indicated on left) in metaphase extract immunodepleted of endogenous CENP-C and M18BP1. Labeling for DNA (Hoechst), total CENP-A, and M18BP1-L (FLAG) is indicated above the image. Scale bar is 5μm. **D**. Quantification of WT or mutant FLAG-M18BP1-L localization with controls (indicated below) in metaphase egg extract immunodepleted of endogenous CENP-C and M18BP1. The CENP-C and M18BP1 depletion and addback condition is indicated in the bottom table. The signal is normalized to WT FLAG-M18BP1-L localization. Error bars represent SEM of three independent replicates (n = 3) with green triangles displaying the mean of each replicate and blue circles representing each individual centromere.

## Discussion

CENP-A nucleosome assembly is decoupled from DNA replication in vertebrates, unlike the assembly of most replication coupled H3 nucleosomes, and instead occurs in G1/interphase (Jansen et al. 2007; Maddox et al. 2007; Moree et al. 2011; Bernad et al. 2011; Silva et al. 2012). In this work, we uncover a new mechanism, wherein the phosphorylation of the conserved site S760 in *X. laevis* M18BP1-L, located in the CENP-C motif, regulates the binding of M18BP1-L to CENP-A nucleosomes and the assembly of new CENP-A nucleosomes in interphase (Fig 5A). The mechanism for phosphorylation disrupting nucleosome binding is likely through the introduction of the negatively charged phosphate group proximal to the positive arginine anchor that engages the acidic patch on H2A/H2B of the nucleosome. This phospho-regulated binding of M18BP1 to CENP-A nucleosomes is a new mechanism that controls the timing of CENP-A assembly.

**Figure 5.**
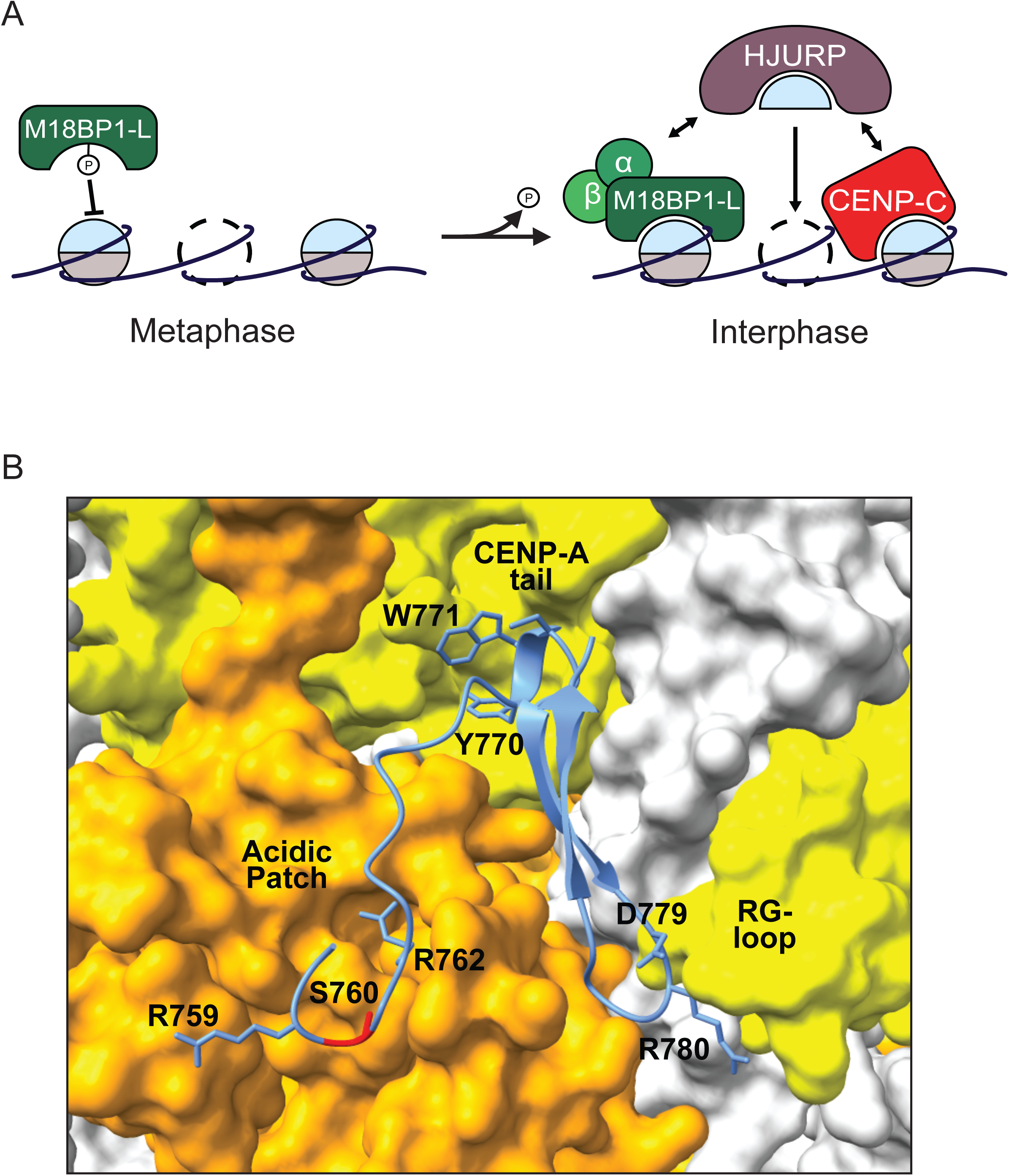
**A**. Model showing phosphorylation dependent inhibition of M18BP1-L binding to CENP-A nucleosomes in metaphase (left) and the dephosphorylation-triggered switch to CENP-A nucleosome binding and new CENP-A assembly in interphase (right). In interphase both M18BP1-L and CENP-C bind to HJURP to recruit new CENP-A for assembly. **B**. AlphaFold structural model of the *X. laevis* CENP-A nucleosome bound to M18BP1-L^758-789^. The surface model of the C-term tail and RG-loop of the CENP-A nucleosome is shown in yellow and the surface model of the acidic patch of H2A/H2B is shown in orange. M18BP1-L is shown in blue, with the conserved residues shown to interact with the CENP-A nucleosome in *G. gallus* labeled (Jiang et al. 2023) and residue S760 highlighted in red.

We tested whether preventing phosphorylation of S760 might be sufficient to drive M18BP1-L binding to CENP-A nucleosomes in metaphase by complementing metaphase extract with S760A mutant M18BP1-L. We showed that the M18BP1-L S760A mutation was unable to bypass the metaphase inhibition of centromere localization, despite our demonstration that M18BP1-L^S760A^ could bind to CENP-A nucleosomes *in vitro*. This indicates that there is another regulatory mechanism preventing M18BP1-L from binding to CENP-A in metaphase. Other post-translational modifications might alter M18BP1-L binding to CENP-A nucleosomes in metaphase. However, other phosphorylatable residues near the regions of M18BP1-L or ggKNL2 that interact with CENP-A were not modified in previous mass spectrometry characterization (French et al. 2017).

An alternative mechanism for preventing M18BP1-L binding to CENP-A in metaphase is that another protein occludes the interaction by binding the CENP-A nucleosome or by binding M18BP1-L. A recent cryo-EM structure of *G. gallus* M18BP1^KNL2^ bound to the CENP-A nucleosome showed that M18BP1^KNL2^ interacts with the C-terminal tail of CENP-A, the acidic patch formed by H2A and H2B, and the RG-loop of CENP-A (Jiang et al. 2023). The sequence of the CENP-A nucleosome binding domain of *X. laevis* M18BP1-L is conserved in *G. gallus* M18BP1^KNL2^ (Fig 1A). We generated an AlphaFold based prediction of the *X. laevis* M18BP1.L bound to the *X. laevis* CENP-A nucleosome which showed that the same contacts are predicted in the *X. laevis* model that were shown to occur in the *G. gallus* structure (Fig 5B). CENP-C engages the same interface on the CENP-A nucleosome (Carroll, Milks, and Straight 2010; Kato et al. 2013; Ariyoshi et al. 2021; Guo et al. 2017) through the CENP-C motif that is conserved in M18BP1. However, we show that depletion of CENP-C does not restore metaphase M18BP1 localization, even in the presence of a non-phosphorylatable S760A mutant, and thus is unlikely to occlude the M18BP1 binding site in metaphase. The only other protein known to bind to CENP-A nucleosomes is CENP-N (Carroll, Milks, and Straight 2010; Carroll et al. 2009). CENP-N interacts with the RG loop of CENP-A nucleosomes (Pentakota et al. 2017; Carroll et al. 2009; Fang et al. 2015; Guo et al. 2017; Carroll, Milks, and Straight 2010; Chittori et al. 2018). As part of the CCAN in metaphase, CENP-N does not bind to CENP-A nucleosomes (Pesenti et al. 2022; Yatskevich et al. 2022; McKinley et al. 2015; Allu et al. 2019). However, an interesting possibility is that CENP-N might bind to CENP-A nucleosomes independent of the CCAN in metaphase. In *G. gallus*, CENP-N and CENP-C compete for binding to CENP-A nucleosomes *in vitro*, and it was proposed that CENP-C increases its affinity for CENP-A nucleosomes in metaphase due to phosphorylation and excludes CENP-N (Ariyoshi et al. 2021). The central domain of *X. laevis* CENP-C lacks the phosphorylation site conserved in humans and *G. gallus* that promotes CENP-A nucleosome binding but retains a threonine at the equivalent residue to S760 in M18BP1 that inhibits nucleosome interaction. Thus it will be interesting to test whether a change in the mode of CENP-C binding might allow CENP-N to engage CENP-A nucleosomes in metaphase and prevent M18BP1-L binding by competing for binding to the RG-loop of CENP-A.

The mechanisms that couple CENP-A assembly to mitotic exit involve a series of phosphorylation dependent protein-protein interactions that inhibit CENP-A assembly until cells complete chromosome segregation and enter G1/interphase. These include multiple steps that alter the activities of the Mis18 complex and CENP-C, the primary adaptors for HJURP and soluble CENP-A recruitment to centromeres. We have discovered an additional control mechanism that prevents CENP-A assembly in metaphase by phosphorylating M18BP1 to prevent the interaction of the CENP-C motif in M18BP1 with the acidic patch of the nucleosome. M18BP1 binds to CENP-A nucleosomes using the C-terminal tail and RG-loop which provide specificity for CENP-A and the acidic patch which increases the affinity of the protein for the nucleosome. Human M18BP1 doesn’t directly bind to the CENP-A nucleosome and also lacks the conserved serine residue that we identified in this work. However, the same residue is conserved in CENP-C motifs across humans, *X. laevis,* and *G. gallus*. It will be interesting to determine whether CENP-C interaction might also be regulated by phosphorylation near the arginine anchor and whether this represents a more generalizable mechanism for tuning the affinity of proteins that bind to centromeric or other nucleosomes.

## Materials and Methods

### Experimental model details

*Xenopus laevis* females (Nasco catalog #LM00535MX) were housed and maintained in the Stanford Veterinary Service Center. Frogs were primed 2 – 14 days prior to ovulation by subcutaneous injection of 50U of Pregnant Mare Serum Gonadotropin (PMSG, BioVendor LLC RP1782725000) in the dorsal lymph sac. Ovulation was induced for 16-18 hrs via subcutaneous injection of 500U Human Chorionic Gonadotropin (HCG, Merck Chorulon Order Code No. CH-475-1) in the dorsal lymph sac. During ovulation, frogs were housed individually in 2L of 1xMMR buffer (6mM Na-HEPES pH 7.8, 0.1mM EDTA, 100mM NaCl, 2mM KCl, 1mM MgCl_2_, 2mM CaCl_2_) at 17°C. All animal work was carried out following the guidelines of the Stanford University Administrative Panel on Laboratory Animal Care (APLAC).

### Protein purification

H2A/H2B dimers and H3/H4 tetramers were purified as described in Guse et al., 2012 and Westhorpe et al., 2015. Histones H2A, H2B, H3, and H4 were expressed from a pST39 vector in BL21-Codon Plus (DE3)-RIPL *Escherichia coli* competent cells and grown in 2xYT media (20g/L tryptone, 10g/L yeast extract, 5g/L NaCl) at 37°C to OD600 of 0.6, then induced with 0.25mM isopropyl β-D-1-thiogalactopyranoside (IPTG) for 3hrs at 37°C. The bacterial cultures were pelleted and resuspended in lysis buffer (20mM Potassium Phosphate (KPO4_4_) pH 6.8, 1M NaCl, 5mM β-mercaptoethanol (β-me), 1mM phenylmethylsulfonyl fluoride (PMSF), 1mM benzamidine, 0.05% NP-40, and 0.2mg/mL lysozyme). The sample was lysed using an EmulsiFlex-C5 (Avestin, Inc.) and 1x 30 sec sonication at duty cycle 50% and output 8 on a Branson Sonicator. The lysed sample was centrifuged at 4°C at 18,000xg for 20 min and the insoluble pellet was washed in lysis buffer, then resuspended in unfolding buffer (7M guanidine-HCl, 20mM Tris-HCl pH 7.5, 10mM DTT). Sample was recentrifuged at 4°C at 18,000xg for 20 min, and the supernatant was dialyzed into urea buffer (6M deionized urea, 200mM NaCl, 10mM Tris-Hcl pH8, 1mM EDTA, 5mM β-me, 0.1mM PMSF). The dialyzed supernatant was then loaded onto 5mL HiTrap Q HP and HiTrap SP HP columns (Cytiva 17115401 and Cytiva 17115201, respectively) in series. Histones were eluted from the Histrap SP column with urea buffer containing 1M NaCl, dialyzed into water, and lyophilized. To reconstitute the H2A/H2B dimer and H3/H4 tetramer, histones were mixed in an equimolar ratio and resuspended in unfolding buffer. They were then dialyzed into 2M NaCl, 10mM Tris-HCl pH 7.6, 1mM EDTA, and 5mM β-me. The samples were then loaded onto a HiLoad 16/600 Superdex 200 column (Cytiva 28989335) and the H2A/H2B dimer and H3/H4 tetramer were purified by size-exclusion chromatography.

*X. laevis* CENP-A-myc/H4 tetramers were expressed and purified as outlined in (Flores Servin, Brown, and Straight 2023) and adapted from (Guse, Fuller, and Straight 2012). CENP-A-myc/H4 was expressed from a pST39 vector in BL21-Codon Plus (DE3)-RIPL *Escherichia coli* competent cells and grown in 2xYT media (20g/L tryptone, 10g/L yeast extract, 5g/L NaCl) at 37°C until OD600 was 0.3-0.4, then moved to 20°C and grown until OD600 was 0.5-0.6. Cultures were then induced for 4hrs with 0.2mM IPTG at 20°C. The bacterial cultures were pelleted and resuspended in lysis buffer (20mM KPO4_4_, 1M NaCl, 5mM β-me, and 1 Roche cOmplete^™^ EDTA-free protease inhibitor cocktail tablet (Millipore Sigma 11873580001). The sample was homogenized using an EmulsiFlex-C5 (3 rounds of 10,000 – 15,000 psi) (Avestin, Inc.) followed by 6x 30 sec sonication at 50% duty cycle and output 8 on a Branson Sonicator. Lysed sample was centrifuged at 4°C for 30 min at 20,000 rpm in a Beckman 45Ti rotor (Beckman Coulter 339160). The supernatant was loaded onto a 30mL hydroxyapatite (HA) column (Type II 20μM HA; Bio-Rad 1572000) pre-equilibrated with 20mM KPO_4_, pH 6.8. The column-bound sample was washed for 6 column volumes (CV) in 20mM KPO_4_ pH 6.8, 1M NaCl, and 5mM β-me, then eluted with 20mM KPO_4_ pH 6.8, 3.5M NaCl, and 5mM β-me. The eluate was dialyzed into S-Column Buffer A (10mM Tris-HCl pH 7.4, 0.75M NaCl, 10mM β-me, and 0.5mM EDTA). The dialyzed protein was loaded onto a 1mL HiTrap SP HP cation exchange column (Cytiva 17115101), washed with 20 CV S-Column Buffer A, followed by a 10 CV wash with 37% S-Column Buffer B (10mM Tris pH 7.4, 0.5mM EDTA, 10mM β-me, 2M NaCl), and eluted over a linear gradient from 37 – 100% S-Column Buffer B. Fractions containing tetramer were pooled, concentrated with an Amicon Ultra Centrifugal Filter (10kDa MWCO; Millipore Sigma UFC801008), aliquoted, flash frozen in liquid nitrogen, and stored at −80°C.

### Chromatin bead reconstitution

Biotinylated 18×601 DNA array was purified as described in Guse et al., 2012. Briefly, puC18 with 18 repeats of the “601” nucleosome positioning sequence (Lowary and Widom 1998) was purified via Gigaprep kit (Qiagen 12191) from STBL2 bacteria grown in Luria Broth (LB) media. The 18×601 plasmid was restriction digested with EcoRI, XbaI, DraI, and HaeII overnight, and purified by polyethylene glycol (PEG) precipitation, using incremental increases of 0.5% PEG (from 4.5 – 10%) with 10 min centrifugation at 5000 x g between each increase. Precipitated and digested 18×601 DNA was dialyzed overnight into TE buffer (10mM Tris, pH 8 and 0.5mM EDTA) and concentrated by ethanol precipitation. The 18×601 DNA was biotinylated by filling EcoRI overhangs with α-thio-dCTP, α-thio-dGTP, α-thio-dTTP (ChemCyte CC-3002-1, CC-3003-1, CC-3004-1), and biotin-14-dATP (Invitrogen 19524016) using Klenow fragment 3’-5’ exo-(New England BioLabs M0212L). Biotinylation of the 18×601 array was verified by combining 500 ng biotinylated 18×601 with 2.5mM NaCl and 1 μg Streptavidin-Alexa 647, and electrophoresing the sample on a 0.7% agarose gel to assay migration.

18×601 reconstituted chromatin was prepared via salt dialysis of 18×601 biotinylated array with purified, recombinant nucleosomes as described in Guse et al., 2012. DNA, H2A/H2B dimer, and either H3/H4 or *xl*CA/H4 tetramer were combined in high-salt buffer (2M NaCl, 10mM Tris pH 7.5, 0.25mM EDTA) and dialyzed over 67 hrs into low-salt buffer (2.5mM NaCl, 10mM Tris pH 7.5, 0.25mM EDTA). H3/H4 or *xl*CA/H4 tetramers were added at a 1.2 x 601 and 1.8 x 601 positioning sequence ratio (respectively) and H2A/H2B dimers were added at a 2.2 x 601 positioning sequence ratio. Reconstituted chromatin was verified by AvaI digestion of the 18×601 chromatin array overnight, followed by SyBr Gold (Life Technologies S11494) staining of a 5% acrylamide native PAGE gel.

Biotinylated 18×601 chromatin array was bound to streptavidin-coated M-280 Dynabeads (Invitrogen 11205D) washed in bead buffer (50mM Tris pH 7.4, 75mM NaCl, 0.25mM EDTA, 0.05% Triton X-100, 2.5% polyvinyl alcohol). Chromatin was bound at 2.6 fmol array per microgram of bead for 60 min at room temp with light shaking. Excess chromatin was removed by multiple washes in bead buffer, and chromatin arrays were incubated in rabbit reticulocyte lysate with *in vitro* translated protein (see below).

### In vitro translation and chromatin binding assays

*In vitro* translated protein was produced in rabbit reticulocyte lysate using the Promega SP6 TNT Quick Coupled Transcription/Translation System (Promega L2080).

To assay binding of *in vitro* transcribed/translated protein to chromatin arrays, rabbit reticulocyte lysate containing the protein of interest was diluted with 5x CSF-XBT (50mM K-KEPES pH 7.7, 500mM KCl, 250mM sucrose, 10mM MgCl_2_, 0.5mM CaCl_2_, 25mM EGTA, 0.25% Triton X-100) to 1x CSF-XBT. 12.5μL of diluted protein-containing lysate was added to 1.5μL chromatin-coated beads and incubated at 21°C for 1hr. Beads were magnetically isolated and washed 3x with 1x CSF-XBT. Beads were boiled for 5 min at 95°C in 10μL protein sample buffer (0.05% bromophenol blue, 10% SDS, 50% glycerol, 200mM Tris pH6.8, 40mM EDTA, 2.86M β-me) and sample was loaded onto a 20% SDS-PAGE gel. Immunoblotting was performed as outlined below.

### *X. laevis* egg extracts

CytoStatic Factor (CSF)-arrested (metaphase) egg extract was produced as described (Desai et al. 1999; Guse, Fuller, and Straight 2012). Following ovulation of the *Xenopus laevis* frogs, eggs were washed 3-5x with MMR buffer (6mM Na-HEPES pH 7.8, 100mM NaCl, 2mM KCl, 2mM CaCl_2_, 1mM MgCl_2_, 0.1mM EDTA) and dejellied in MMR + 2% (w/v) L-cysteine for 5min. De-jellied eggs were washed 3-5x in 1x CSF-XB buffer (10mM K-HEPES pH 7.7, 100mM KCl, 50mM sucrose, 2mM MgCl_2_, 0.1mM CaCl_2_, 5mM K-EGTA), then washed into 1x CSF-XB buffer + 10 μg/mL LPC (Leupeptin/Pepstatin A/Chemostatin). Eggs were packed in a 13 x 51mm polyallomer tube (Beckman Coulter, 326819) by centrifugation in an International Clinical Centrifuge (model CL no. 740250; rotor model 221) at ∼1250 rpm for 30s then ∼1900 rpm for 15s at room temperature. Excess buffer was removed and packed eggs were crushed by centrifugation in a SW55Ti rotor (Beckman Coulter 342196) for 15 min at 10,000 rpm (16°C). The soluble cytoplasmic fraction was removed by side-puncture of the tube with a 16G, 1.5in needle and supplemented with 10 μg/ml LPC, 10 μg/ml cytochalasin D, 50 mM sucrose, and energy mix (7.5 mM creatine phosphate, 1 mM ATP, and 1 mM MgCl_2_).

### Immunodepletions

Immunodepletion of endogenous M18BP1 and CENP-C from *Xenopus laevis* egg extract was performed as described (Moree et al. 2011). Protein A Dynabeads (Invitrogen, 10001D) were washed in 3x CSF-XBT (10mM K-HEPES pH 7.7, 100mM KCl, 50mM sucrose, 2mM MgCl_2_, 0.1mM CaCl_2_, 5mM K-EGTA, 0.05% Triton X-100) and coupled to antibody for 60 min at 4°C on a rotator. To immunodeplete CENP-C from 100μL of egg extract, 5μg of anti-xlCENP-C antibody (rabbit, raised and purified against xlCENP-C^207-296^ (Milks, Moree, and Straight 2009))) was bound to 33μL of Protein A Dynabeads. To immunodeplete M18BP1 from 100μL of egg extract, 5μg of anti-xlM18BP1 (rabbit, raised against GST-xlM18BP1-L^161-415^ and purified against xlM18BP1.S^161-375^) was bound to 33μL of Protein A Dynabeads. For the mock depletion of 100 μL of egg extract, 5μg of whole rabbit IgG (Jackson ImmunoResearch Laboratories, Inc. 011-000-003) was bound to 33μL of Protein A Dynabeads. After coupling to antibody, beads were washed 3x in CSF-XBT and resuspended in egg extract. The extract was depleted for 1hr at 4°C on a rotator, and the beads were removed by 3x 5 min magnetic pull-downs. Immunodepletion of the egg extract was verified by western blotting.

### CENP-A assembly assays

CENP-A assembly on sperm chromatin was performed as described (Moree et al. 2011; Westhorpe, Fuller, and Straight 2015). V5-CENP-A mRNA and HJURP mRNA for use in egg extracts was produced using the Invitrogen mMessage mMachine SP6 Transcription kit (Fisher Scientific, AM1340) following the manufacturer instructions, except 3μg of pCS2+ plasmid was linearized via NotI digestion for the transcription reaction. Following transcription, mRNA was purified using RNeasy mini columns (Qiagen 74104) according to manufacturer instructions.

CENP-A assembly on sperm chromatin was assayed by setting up 30μL assembly reactions containing 25ng/μL V5-CENP-A mRNA, 40ng/μL HJURP mRNA, 3μL FLAG-M18BP1-L IVT protein, and 25μL of M18BP1 and/or CENP-C immunodepleted, CSF-arrested egg extract. RNA and IVT-supplemented egg extract was incubated at 16-18°C for 30min (flicking tubes every 15min) to allow RNA translation to occur. The translation reaction was terminated by the addition of 0.1mg/ml cycloheximide and reactions were released into interphase by the addition of 750μM CaCl_2_. Egg extract reactions were supplemented with 3000 demembranated sperm chromatin per μL. Reactions were incubated at 16-18°C for 75min, then diluted into 1mL dilution buffer (BRB-80 [80mM K-PIPES pH 6.8, 1mM MgCl_2_, 1mM EGTA], 150mM KCl, 0.5% Triton X-100, 30% glycerol) and incubated on ice for 5min. To each reaction, 1mL of fixation buffer (dilution buffer + 4% formaldehyde) was added, and reactions were layered onto a cushion of BRB-80 + 40% glycerol in 15mL glass centrifuge tubes (Kimble 45500-15). Sperm nuclei were spun down onto acid-washed, poly-L-lysine-coated coverslips via centrifugation at 3500rpm for 20min and then further processed for immunofluorescence.

### Immunofluorescence preparation

Coverslips were blocked in Antibody Dilution Buffer (AbDil) (20 mM Tris-HCl, pH 7.4, 150 mM NaCl with 0.1% Triton X-100, and 2% bovine serum albumin) for 30min. Coverslips were exposed to primary antibody diluted in AbDil for 30min (1μg/mL rabbit anti-xlCENP-A (raised and purified against xlCENP-A^1-50^), 5μg/mL mouse anti-FLAG (Millipore Sigma F1804), and/or 2μg/mL mouse anti-V5 (ThermoFisher R960-25)). Coverslips were washed 3x in AbDil, then exposed to secondary antibody diluted in AbDil for 30min (1.5μg/mL Alexa 647 goat anti-mouse IgG (Invitrogen A21236) and 1.5μg/mL Alexa 568 goat anti-rabbit IgG (Invitrogen A11011).

Coverslips were washed 3x in AbDil, then stained for 5min with 10μg/mL hoechst in AbDil (ThermoFisher H3570) for DNA, and washed 2x in AbDil and 2x in 1x Phosphate-Buffered Saline (PBS). Coverslips were mounted with mounting media (90% glycerol, 10 mM Tris-HCl, pH 8.8, 0.5% p-phenylenediamine) and sealed with clear nail polish and stored at –20°C.

### Image acquisition and processing

Imaging was performed on a Nikon Eclipse Ti2 inverted microscope (Nikon TI2-LA-FL-2) with a CrestOptic X-Light V3 Confocal Unit (Nikon) and a Celesta Light Engine (Lumencor), controlled via NIS-Elements 6.10.01 software (Nikon). Images of sperm nuclei were acquired with a Nikon Plan Apo 60XA/1.40 oil immersion lens. Images were acquired using an Electron Multiplying Charge-Coupled Device (EMCCD) camera (iXon Life 897) and digitized to 16 bits. Z sections were taken at 0.2μm intervals across a span of 8μm. Displayed images of sperm nuclei are maximum intensity projections of z-stacks with brightness adjusted uniformly within a figure to facilitate viewing.

### Immunoblotting

Samples were resolved by SDS-PAGE and transferred onto nitrocellulose membrane (Amersham GE10-6000-00). All samples were transferred in CAPS transfer buffer (10mM 3-(cyclohexylamino)-1-propanesulfonic acid, pH 11.3; 0.1% SDS; 20% methanol). Samples containing 0.1μL IVT protein and 1uL *Xenopus* egg extract were loaded per lane.

Membranes were blocked in 4% dry milk in TBSTx (20mM Tris, 150mM NaCl, 0.1% Tween-20) and probed with primary antibody. Primary antibodies used were 2μg/mL rabbit-anti-xlM18BP1 (raised against GST-xlM18BP1-L^161–415^ and purified against xlM18BP1-S^161–375^ (Moree et al. 2011), 1.5μg/mL rabbit-anti-xlCENP-C (raised and purified against xCENP-C^207–296^ (Milks, Moree, and Straight 2009), 1.5μg/mL mouse-anti-DM1α (tubulin; Sigma T6199), and 1μg/mL mouse-anti-myc clone 4A6 (Sigma 05-724MG). Detection was performed using horseradish peroxidase-conjugated goat-anti-mouse or goat-anti-rabbit secondary antibodies (Bio-Rad 1706516, 1706515) followed by chemiluminescence (Pierce ECL Western Blotting Substrate, Thermofisher 32106). Chemiluminescent membranes were exposed to X-ray film for 30-300sec and developed with a Konica SRX-101A film processor.

### Protein alignment

Protein sequences homologous to *X. laevis* M18BP1-L^758-779^ were identified using NCBI PSI-BLAST. Multiple sequence alignment was performed in Snapgene using the MAFFT algorithm (Katoh et al. 2002).

### Quantification and statistical analysis

Image analysis of *Xenopus laevis* sperm was performed using custom software as described (Moree et al. 2011) and publicly available at https://github.com/cjfuller/imageanalysistools. To identify centromeres, images were normalized by median filtering the image and dividing the original image by the filtered intensity value. The channel was thresholded with the indicated centromere marker (generally CENP-A) to generate a centromere mask, and then filtered by size to remove regions larger or smaller than centromeres (using a minimum cut-off of 5 pixels and maximum cut-off of 25 pixels). Masks were then manually inspected to remove remaining non-centromere regions. Mean pixel intensity per centromere region was measured for channels of interest using maximum intensity projections. Values for each centromere region are plotted in each dot plot as circles for every experiment and condition and normalized to the WT, with the mean of each biological replicate shown as triangles, and the overall mean and standard error shown with lines. For sperm, at least 100 centromeres among at least 10 nuclei were counted for each condition in each replicate in each experiment. All M18BP1 localization experiments were repeated for 3 biological replicates (n = 3) and all CENP-A assembly experiments were repeated for 4 biological replicates (n = 4).

Western blots were quantified using ImageJ (National Institutes of Health). A rectangle of fixed size was drawn around every band (both bound protein and loading control) and mean grey value was measured. Background was calculated by taking the mean grey value for a region of the same size below each band. Background value was subtracted from the values measured for each band, then each band was first normalized to its respective loading control, and finally normalized to an average of the values for the bands representing WT M18BP1 binding to CENP-A.

All plotting and statistics were performed using GraphPad Prism. Every relevant experiment was analyzed using a Welch’s unpaired two-tailed t-test on the means of the n = 3 or n = 4 biological replicates.

## Acknowledgements

The authors are thankful to Pragya Sidhwani for feedback on the manuscript. This work was supported by the grant of National Institutes of Health R01 GM074728 to Aaron Straight; Jacob Price Schwartz and Rae Brown were supported by National Institutes of Health 5 T32 GM007276; and Rae Brown was supported by a National Science Foundation Graduate Research Fellowship DGE-1656518.

## Author contributions

**Rae Brown**: Conceptualization, Methodology, Investigation, Data Curation, Writing, Visualization; **Jacob Price Schwartz**: Methodology, Investigation; **Lyin Ghadri**: Investigation; **Aaron Straight**: Conceptualization, Methodology, Supervision, Writing, Visualization, Project Administration, Funding Acquisition

## Synopsis

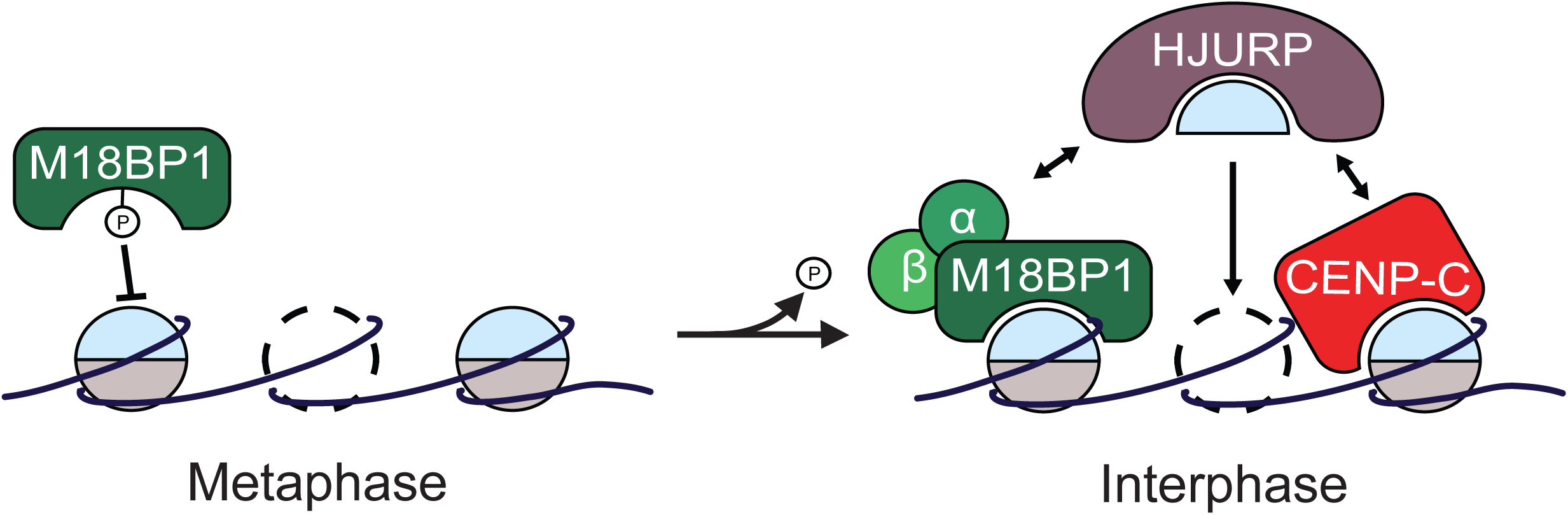

**Figure EV1.**
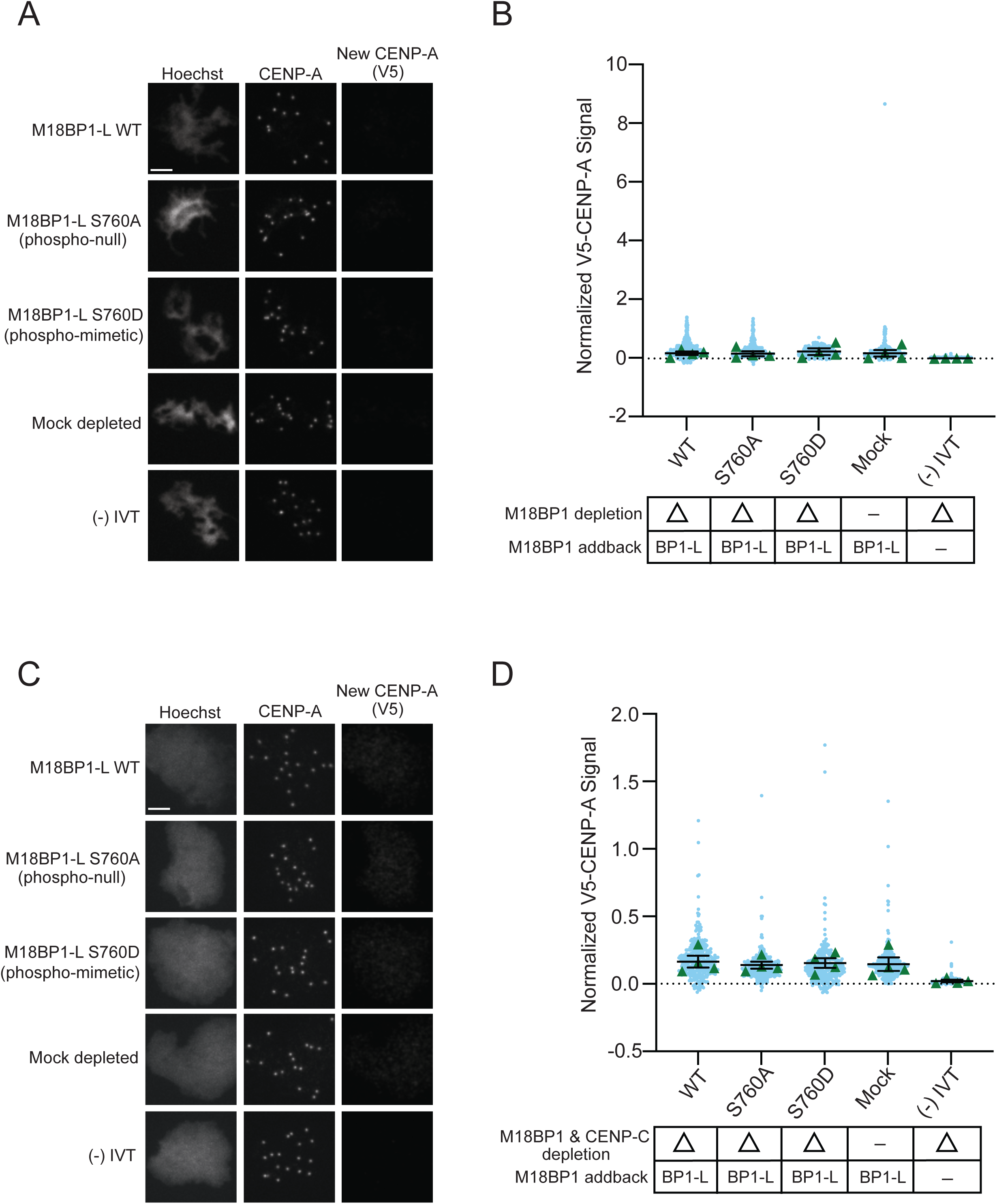
**A**. Representative immunofluorescence images of new V5-CENP-A assembly in metaphase extract immunodepleted of endogenous M18BP1 then supplemented with full-length WT or mutant FLAG-M18BP1-L or a mock depletion or (-) IVT control (indicated on left). Labeling for DNA (Hoechst), total CENP-A, and new CENP-A (V5) is indicated above the image. Scale bar is 5μm. **B**. Quantification of new V5-CENP-A with controls (indicated below) in metaphase egg extract immunodepleted of endogenous M18BP1. M18BP1 depletion and addback condition is indicated in the bottom table. The signal is normalized to the WT FLAG-M18BP1-L addback condition. Error bars represent SEM of four independent replicates (n = 4) with green triangles displaying the mean of each replicate and blue circles representing each individual centromere. **C**. Representative immunofluorescence images of new V5-CENP-A assembly in metaphase extract immunodepleted of endogenous CENP-C and M18BP1 then supplemented with full-length WT or mutant FLAG-M18BP1-L or a mock depletion or (-) IVT control (indicated on left). Labeling for DNA (Hoechst), total CENP-A, and new CENP-A (V5) is indicated above the image. Scale bar is 5μm. **D**. Quantification of new V5-CENP-A with controls (indicated below) in metaphase egg extract immunodepleted of endogenous CENP-C and M18BP1. CENP-C and M18BP1 depletion and addback condition is indicated in the bottom table. The signal is normalized to the WT FLAG-M18BP1-L addback condition. Error bars represent SEM of four independent replicates (n = 4) with green triangles displaying the mean of each replicate and blue circles representing each individual centromere.

